# Graph attention-based fusion of pathology images and gene expression for prediction of cancer survival

**DOI:** 10.1101/2023.10.26.564236

**Authors:** Yi Zheng, Regan D. Conrad, Emily J. Green, Eric J. Burks, Margrit Betke, Jennifer E. Beane, Vijaya B. Kolachalama

## Abstract

Multimodal machine learning models are being developed to analyze pathology images and other modalities, such as gene expression, to gain clinical and biological insights. However, most frameworks for multimodal data fusion do not fully account for the interactions between different modalities. Here, we present an attention-based fusion architecture that integrates a graph representation of pathology images with gene expression data and concomitantly learns from the fused information to predict patient-specific survival. In our approach, pathology images are represented as undirected graphs, and their embeddings are combined with embeddings of gene expression signatures using an attention mechanism to stratify tumors by patient survival. We show that our framework improves the survival prediction of human non-small cell lung cancers, out-performing existing state-of-the-art approaches that leverage multimodal data. Our framework can facilitate spatial molecular profiling to identify tumor heterogeneity using pathology images and gene expression data, complementing results obtained from more expensive spatial transcriptomic and proteomic technologies.

## 1 Introduction

The field of spatial biology is rapidly expanding as technologies such as spatial proteomics and transcriptomics seek to unravel the complex spatial organization of cells and how it influences cellular phenotypes in health and disease. The three-dimensional organization of cells into tissue microenvironments has a significant impact on disease development, progression, and outcomes. Spatial omic technologies are also enabling connections between single cell omic profiles and pathology. Many spatial technologies produce spatial omic data on the same tissue specimen stained using hematoxylin and eosin (H&E) and digitized to produce a standard pathology whole slide image (WSI). Several methods combining spatial omic data and extracted pathology features have been published [10, 25, 31] to improve cell type identification, cell type deconvolution, spatial pattern recognition, and predict omic features on pathology images alone. While these technologies and methods are promising, the data are expensive and technically challenging to generate, resulting in a small number of cases profiled that may only capture a portion of the entire WSI. We sought to develop a method to utilize digitized H&E WSIs and bulk-derived omic data, which are less expensive to collect and often present across large number of samples, to spatially localize predictive features and characterize disease-associated alterations in tissue microenvironments. The method allows utilization of large public resources of WSIs and bulk omic data, such as The Cancer Genome Atlas (TCGA) [26], to identify interesting spatially resolved disease-associated alterations. The method can be used to generate hypotheses and identify regions of interest within tissues based on large sample sets that can be further characterized using modern spatial technologies.

Digitized H&E WSIs have been used in advanced machine learning frameworks, computer vision, and multimodal learning to quantify the molecular underpinnings of disease, estimate markers of disease progression, and predict patient survival. Computer methods to analyze WSIs for automated diagnosis and quantification of morphologic biomarkers have seen remarkable progress. Methods have been developed to analyze multimodal datasets that predict outcome metrics such as survival by combining clinical, imaging, and genomic data using fusion frameworks. Chen and colleagues recently developed a weakly-supervised, late-fusion framework to combine WSIs and corresponding bulk genomic data and predict survival on various cancers [3]. However, learning the spatial relevance of non-imaging data such as bulk gene expression is not straightforward using late fusion. To understand the spatial relationships governing disease-associated alterations in the tissue microenvironment, we sought an approach that integrates digitized WSIs and bulk omic data and learns early in the data fusion training cycle. While other researchers have previously explored the development of mid-level-fusion and mixer architectures [6, 28] as well as the use of graph-based representations of WSIs [37], our work is unique in mixing node and edge embeddings along with fusion of bulk gene expression embeddings to learn a multimodal topographic mapping to predict survival.

Our framework (Fig. 1), allows for representation of WSIs in the form of undirected graphs (Fig. 2), whose embeddings are fused with embeddings of bulk omic data to predict patient survival. In the graph, nodes represent local image patches and edges represent patch adjacency. Using the WSI-graph as input, we define a graph-mixer module that comprises of node-mixing and channel-mixing layers for learning relationships between neighboring nodes and representative features of each node on the WSI-graph, respectively. We pass the resulting embeddings into an attention module, which also receives embeddings of genomic signatures as input, and thus captures local interactions between image and genomic data. Our framework then passes the image-genomic embeddings to a global attention pooling layer and a subsequent fully-connected layer to predict survival risk. The spatially-resolved multimodal features that our framework computes in this fashion can be used to understand changes in the tissue microenvironment that are predictive of patient survival. Our experiments show that our framework is highly adaptable and can be used on a variety of bulk omic datasets and corresponding WSIs in various disease contexts.

**Figure 1.**
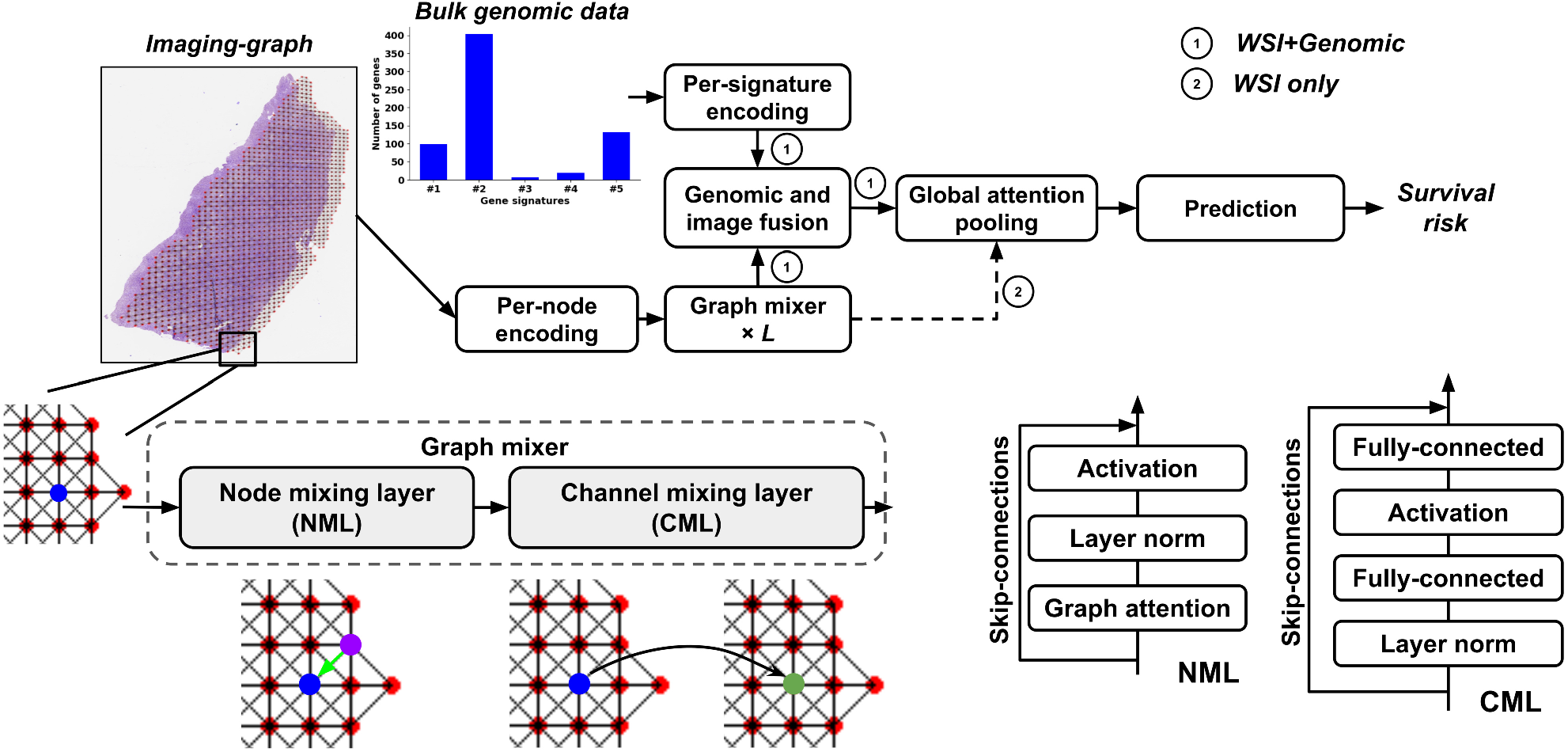
Graph attention-based fusion framework. The mixer framework (left) uses the graph node embeddings and gene expression signature embeddings and jointly learns a spatial fingerprint of the WSI-transcriptomic relationship via an attention-based framework to predict survival. The graph mixer (right) comprises of consecutive node-mixing and channel-mixing layers for learning relationships between adjacent (blue and purple) nodes and more representative node features (blue to green) on the graph. The per-node encoding, per-signature encoding, and prediction modules consist of fully-connected layers. The details of graph mixer, genomic and image fusion, and global attention pooling modules are described in Section 2.2.

**Figure 2.**
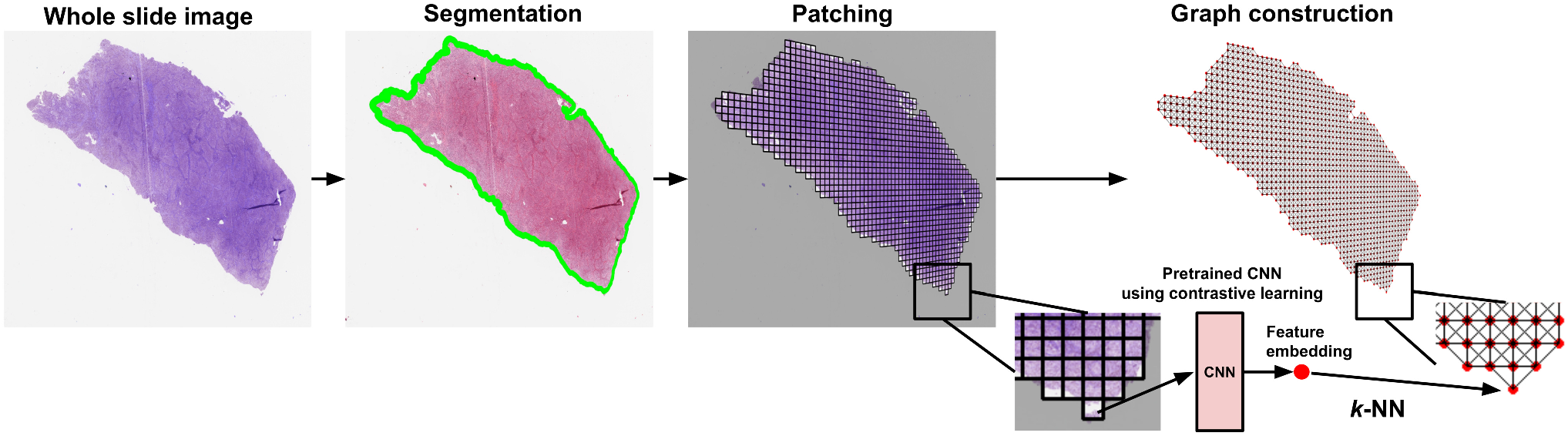
Whole slide image (WSI) processing and graph construction. WSIs were processed using a pipeline involving foreground-background separation, tessellation into image patches followed by construction of an undirected graph. Patch embeddings were generated using a contrastive learning framework (Fig. 3) and used as node features in the graph.

### 1.1 Contributions

We summarize the key contributions of this work as follows:

- We developed a multimodal data fusion architecture that combines embeddings of WSIs, represented as undirected graphs, with embeddings of gene expression signatures using an attention-based mechanism to predict patient survival. Our architecture is unique in its graph-based modeling of local and global features as well as interpretation of image-genomic interactions.
- Our experiments show that our framework achieves state-of-the-art performance in predicting survival on human non-small cell lung cancers (NSCLC): lung adenocarcinoma (LUAD) and lung squamous cell carcinoma (LUSC), which are the two most common histological types of NSCLCs.
- We introduce survival activation maps (SAM), which are saliency-based spatial signatures on WSIs that highlight tumor regions associated with the output of interest. SAM can incorporate gene expression-specific information on WSIs and generate multimodal spatial signatures that may provide insights into tissue features associated with patient survival.

## 2 Materials and methods

### 2.1 Study population

We obtained WSIs, bulk gene expression data, demographic, and clinical (including overall survival) data on subjects with LUAD or LUSC from The Cancer Genome Atlas (TCGA) [26], the Clinical Proteomic Tumor Analysis Consortium (CPTAC) [8], and the National Lung Screening Trial (NLST) [27] (Table 1). TCGA is a landmark cancer genomics program that characterized molecular alterations in thousands of primary cancer and matched normal samples spanning several cancer types. CPTAC is a national effort to accelerate the understanding of the molecular basis of cancer through the application of large-scale proteome and genome analysis. NLST was a randomized controlled trial to determine whether screening for lung cancer with low-dose helical computed tomography reduces mortality from lung cancer in high-risk individuals relative to screening with chest radiography.

**Table 1:** Study population. Source: Online portals of the TCGA, CPTAC, and NLST cohorts.

| Dataset<br>[subjects] | Age mean<br>(std) | Gender male<br>(percent) | Uncensored<br>(percent) | Survival time in days<br>[min, max] (median) | <sup>1</sup>Race<br>information | <sup>2</sup>Stage<br>information |
| --- | --- | --- | --- | --- | --- | --- |
| TCGA LUAD [n=444] | 65 (10.0) | 203 (45.72%) | 156 (35.14%) | [4, 7143] (658) | (342, 51, 7, 1, 43) | (245, 109, 52, 23, 15) |
| TCGA LUSC [n=471] | 67 (8.5) | 352 (74.73%) | 202 (42.89%) | [0, 4765] (641) | (324, 30, 9, 0, 108) | (230, 151, 62, 6, 22) |
| CPTAC LUAD [n=199] | 63 (9.3) | 129 (64.82%) | 35 (17.59%) | [0, 1836] (456) | (53, 4, 1, 1, 140) | (104, 49, 42, 2, 2) |
| CPTAC LUSC [n=102] | 66 (8.4) | 83 (81.37%) | 19 (18.62%) | [0, 1785] (742) | (30, 0, 0, 0, 72) | (37, 44, 19, 1, 1) |
| <sup>3</sup>NLST LUAD [n=229] | 64 (5.2) | 122 (53.28%) | 68 (29.69%) | [189, 2786] (2425) | (213, 8, 5, 1, 2) | (159, 24, 33, 13, 0) |
| <sup>3</sup>NLST LUSC [n=115] | 64 (4.9) | 84 (73.04%) | 38 (33.04%) | [328, 2751] (2379) | (102, 5, 6, 0, 2) | (84, 13, 14, 4, 0) |

<sup>1</sup> White; Black; Asian; American Indian or Alaska Native; Unknown.
<sup>2</sup> Stage I; Stage II; Stage III; Stage IV; Unknown.
<sup>3</sup> NLST is only used for fine-tuning feature generation in Fig. 3.

Several studies have reported gene expression signatures associated with lung cancer survival. To demonstrate a proof-of-concept, we focused on gene signatures associated with B cell populations. B cell associated signatures have been shown to be elevated in both LUAD and LUSC; however, increased tumor-infiltrating B cells are associated with good prognosis only in LUAD. We included 5 gene expression signatures specific for B cell populations derived from single-cell RNA sequencing data profiling of normal adjacent lung tissue and lung cancer tissue: *Sinjab (Plasma)*, denoted as sig#1, *Sinjab (B Cell)*, denoted as sig#2, *Sinjab (B:1)*, denoted as sig#3, *Sinjab (B: 0)*, denoted as sig#4 [22], and *Travaglini (B)*, denoted as sig#5 [29].

### 2.2 Modeling framework

Our framework jointly learns to interpret WSIs and corresponding genomic data to predict tumor survival, and generates spatial image-genomic signatures that point to tumor regions that are highly associated with patient survival. We developed two survival models: (a) WSI-only model denoted as imaging survival model (ISM), and (b) model that integrates WSIs and genomic data, denoted as fusion survival model (FSM).

#### 2.2.1 Whole slide image processing and graph construction

Let *G* = (*V, E*) be an undirected graph where *V* is the set of nodes representing the image patches of the WSI and *E* is the set of edges between the nodes in *V* that represent whether two image patches are adjacent to each other (Fig. 2). We denote the adjacency matrix of *G* as A = [A_*ij*_] where A_*ij*_ = 1 if there exists an edge (*v*_*i*_, *v*_*j*_) ∈ *E* and A_*ij*_ = 0 otherwise. An image patch must be connected to other patches and can be surrounded by at most 8 adjacent patches, so the sum of each row or column of *A* is at least one and at most 8. A graph can be associated with a node feature matrix *H, H* ∈ IR^*N×C*^, where each row contains the *C*-dimensional feature vector computed for an image patch, i.e., node, and *N* = |*V*|. The C-dimensional feature vector is obtained by passing an image patch through a convolutional neural network (CNN) that has been trained using contrastive learning [7] (Fig. 3). We refer to the graph representation of the WSI as the imaging-graph, *IG* = (*H, A*).

**Figure 3.**
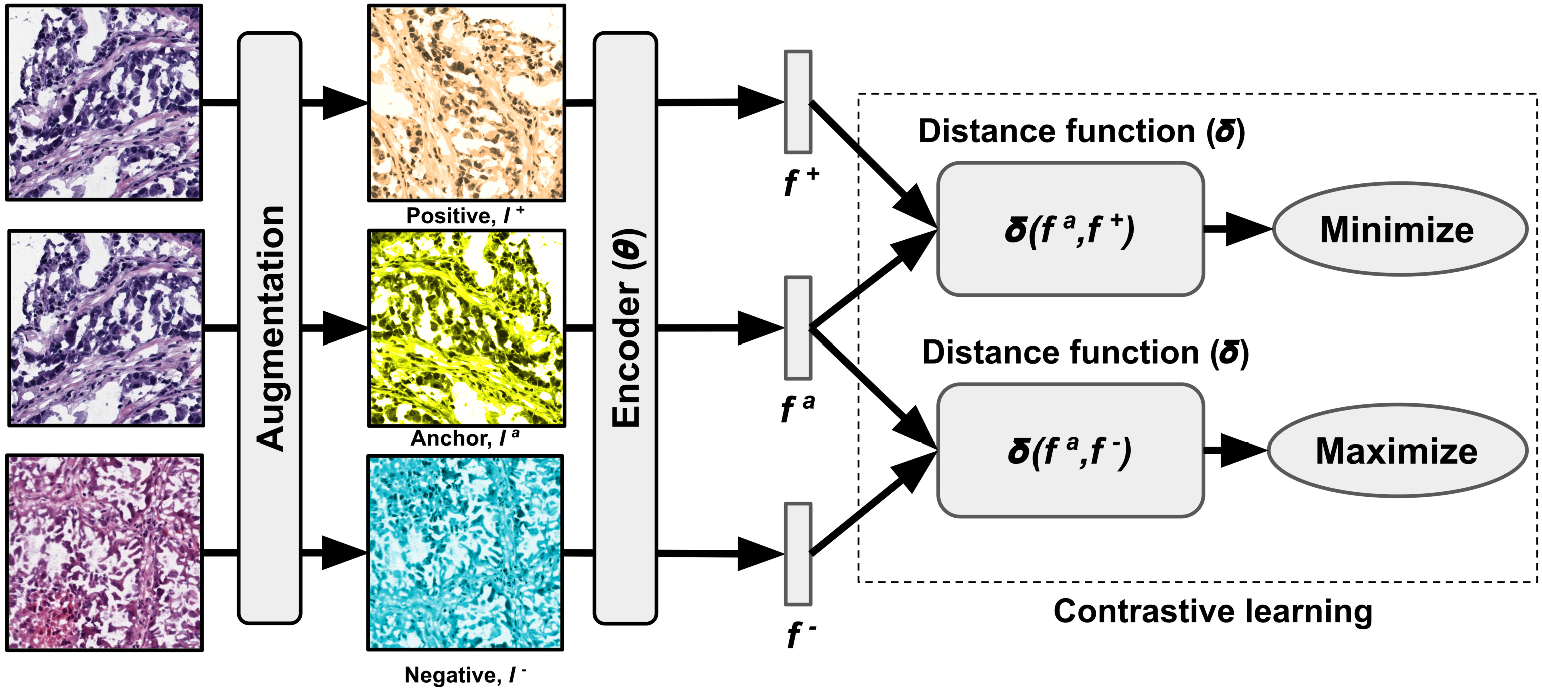
Feature generation and contrastive learning. We applied three distinct augmentation functions, including random color distortions, random Gaussian blur, and random cropping followed by resizing back to the original size. The encoder, *θ*, received an augmented image and generates an embedding vector as the output. These vectors were used for computing contrastive learning loss to train the encoder. After training, we used the embedding vectors for graph construction.

#### 2.2.2 Node and channel mixing

Our framework is built using the imaging graph *IG* mixed with corresponding bulk gene expression data. It consists of a per-node embedding layer, a stack of *L* identical Graph-Mixer layers, *M* per-signature encoding layers, a genomic and image fusion module, a global attention-pooling layer, and a fully-connected layer as the final prediction layer. Our framework without per-signature encoding layers and the genomic and image fusion module can work on WSIs as the only input, and we refer to this model as imaging survival model (ISM)). The core Graph-Mixer layer has two parts: a node mixing layer (NML) and a channel mixing layer (CML).

The input graph node embeddings were mapped to latent space via the per-node embedding module, where *H* ∈ IR^*N×C*^ →*H* ∈ IR^*N×D*^ and *D* is the hidden size. The well-known MLP-Mixer [28] works only on a fixed number of tokens and becomes less effective in handling graph-structured data. Given that the number of nodes in *G* across all WSIs is variable, we addressed this via our architecture, which resembles the GraphMLP framework [17], recently proposed for human pose estimation. This framework learns local and global information of the imaging-graph. In contrast to GraphMLP, we applied the graph attention layer just on the node mixing layer for token mixing [28]. The graph attention layer makes every node in *G* attend to its neighbors given its own representation as the query, so that the local relationships are better learned than the MLP-Mixer or the GraphMLP.

Specifically, our GraphMixer layer is composed of a node mixing layer (NML) and a channel mixing layer (CML) (Fig. 1). The NML contains a graph attention layer, which is built upon Graph Attention Network (GAT) [2]. Unlike the Graph Convolution Network (GCN) used in GraphMLP, which weighs all neighbors *N*_*i*_ for a given node *i* in *G* with equal importance, GAT computes a learned weighted average of the representations of *N*_*i*_. It computes a score for every edge (*j, i*), which indicates the importance of the features of the neighbor *j* to the node *i*:

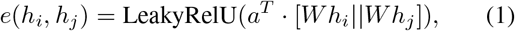

where *a* ∈ IR^2*D*^, *W*∈ IR^*D×D*^ are learned, and || denotes vector concatenation. These attention scores are normalized across all neighbors *j ∈ N*_*i*_ using softmax, and the attention function is defined as:

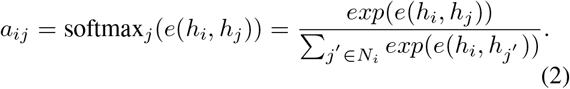

We then computed a weighted average of the transformed features of the neighbor nodes (followed by a nonlinearity *σ*) as the new representation of node *i*, using the normalized attention coefficients:

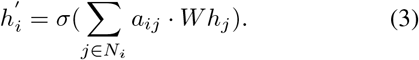

We refer to the previous three equations as the GA(·). The CML has a similar architecture to MLP-Mixer with the channel MLP and has no matrix transposition. Based on the above description, the GraphMixer layer processes image-graph *IG* = (*H, A*) as:

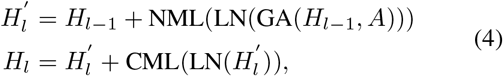

where *l*∈ [1, …, *L*] is the index of GraphMixer layers. Here 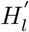 and *H*_*l*_ are the output features of the NML and the CML for GraphMixer layer *l*, respectively.

#### 2.2.3 Genomic signature embeddings

Gene counts derived from bulk RNA sequencing data from LUAD (229 CPTAC; 517 TCGA) and LUSC (109 CPTAC; 501 TCGA) tumor samples were obtained from the Genomic Data Commons [9]. For each dataset (CPTAC-LUAD, TCGA-LUAD, CPTAC-LUSC, TCGA-LUSC), duplicate samples and low-signal or invariant genes were filtered out. Specifically, gene filtering was conducted on normalized gene count data (the EdgeR Bioconductor package was used to compute log2 counts per million using library sizes estimated using the trimmed mean of M-values method) [19], by removing genes with a zero interquartile range or a cumulative sum across samples equal to or below one. Duplicate gene names were collapsed using WGCNA’s ‘collapseRows’ function with the default ‘maxMean’ method [16]. The final set of genes (n=12,306 genes) was the union set of LUAD genes (n=11,975 intersecting genes between TCGA-LUAD and CPTAC-LUAD) and LUSC genes (n=11,933 intersecting genes between TCGA-LUSC and CPTAC-LUSC). Each dataset was renormalized as described above using the final set of genes.

Batch correction was performed separately for LUAD and LUSC samples using ComBat [13], with TCGA serving as the reference batch for both. Using the batch corrected and normalized gene matrices for LUAD and LUSC, we encoded each gene signature into embeddings using a fully-connected layer to get feature representations. Let 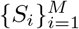 be *M* unique gene signatures associated with distinct biological functions or clinical phenotypes (e.g., overall survival), where *S*_*i*_ ∈ IR^*P ×*1^ with *P* genes and *P* is variant for different signatures. We used the trainable per-signature encoding layer to encode *S*_*i*_ to a D-dimensional genomic signature embedding *B*_*i*_ = Φ_*i*_(*S*_*i*_), where *B*_*i*_ ∈ IR^*D×*1^. Finally, we concatenated all *M* signature embeddings *B*_*i*_ together as *B*, where *B* ∈ IR^*M×D*^.

#### 2.2.4 Genomic and image fusion

We leveraged a Query-Key-Value (QKV) mechanism to capture interpretable image-genomic interactions that exist in the tumor microenvironment (Fig. 1). This framework was inspired by prior work [6, 30], which directly models pairwise interactions between each node in *IG* and each genomic signature. We denote this approach to genomic and image fusion as the Genomic Attention Module (GAM). The GAM attention uses genomic signature embeddings to encode the imaging-graph features into imaging-genomic features, using the following mapping:

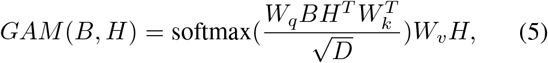

where *W*_*q*_, *W*_*k*_, *W*_*v*_ ∈ IR^*D×D*^ are trainable weights, *B* ∈ IR^*M×D*^ are the genomic signature embeddings, and *H* ∈ IR^*N×D*^ are the nodes embeddings after *L* GraphMixer layers.

#### 2.2.5 Global attention pooling

Inspired by [12], we proposed a gating-based weighted average of nodes where weights are determined by a neural network. Additionally, the weights must sum to 1 to be invariant to the size of *IG*. Let *H* = {*h*_1_, …, *h*_*N*_} be node features after the *L* GraphMixer layers (*L* = 3 in our case), and we propose the following global attention pooling:

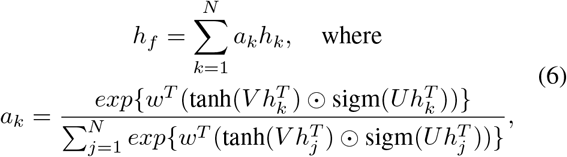

*w* ∈ IR^*L×*1^ and *V, U* ∈ IR^*L×M*^ are learnable parameters, ⊙ is an element-wise multiplication and sigm(·) is the sigmoid non-linearity.

#### 2.2.6 Survival loss function

The pooled WSI-level embedding after global attention pooling was subsequently supervised using the cross entropy-based Cox proportional loss function for survival analysis [3]. We first partitioned the continuous timescale of overall patient survival time in months, *T* into 4 non-overlapping bins: [*t*_0_, *t*_1_), [*t*_1_, *t*_2_), [*t*_2_, *t*_3_), [*t*_3_, *t*_4_), where *t*_0_ = 0, *t*_4_ = ∞ and *t*_1_, *t*_2_, *t*_3_ define the quartiles of event times for uncensored patients in the TCGA cohort. The discretized event time *Y*_*i*_ of patient *i* with continuous event time *T*_*i*_ is then defined as:

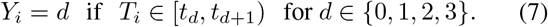

For a given patient *i* with the discrete event time *Y*_*i*_ and *h*_*f*_ after global attention pooling, we modeled the hazard function using the sigmoid activation defined as:

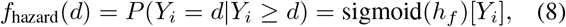

where [*Y*_*i*_] means getting value of index *Y*_*i*_, and the survival function is then defined as:

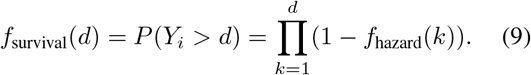

The loss *L* during the training is defined using the log-likelihood function for a discrete survival model [36] as *N* → *M* → 1:

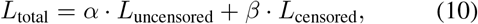

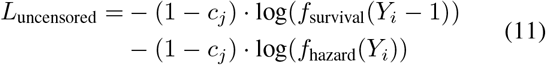

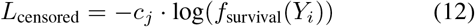

### 2.3 Model interpretability

Interpretability methods, such as GradCAM [20], provide valuable visual perspectives on the inner workings of neural networks, especially in the context of image classification. Specifically tailored for convolutional neural networks (CNNs), GradCAM typically concentrates on the final convolutional layer, emphasizing the significant regions of an image that influence class predictions. Nevertheless, the dimensions of this layer mean the derived heatmap is inherently of a coarse resolution. Consequently, GradCAM does not directly align with our model’s structure, which does not utilize convolutional layers.

We adapted the GradCAM framework to address the aforementioned challenges and identify spatial features that are highly associated with tumor survival. We denoted the interpretations as survival activation maps (SAM). First, we computed the gradients of logits of the *h*_*f*_ for the first survival time bin, with respect to feature maps *A*_*j*_ of the last GraphMixer layer. This approach would produce fine-grained localization maps compared to GradCAM. It is because GradCAM depends on the feature maps from layers that have been subjected to pooling, potentially losing detailed spatial information. Then these gradients flowing back are average-pooled over *N* nodes in the imaging-graph to obtain the importance weights *α*_*j*_ for each feature map *A*_*j*_:

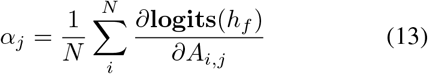

We computed the weighted sum of the feature maps using the importance weights *α*_*j*_ to obtain the visualization of the areas that contributed most to tumor survival:

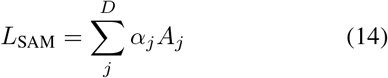

GradCAM typically highlights areas in the image that positively contribute to a class. It might not clearly show regions that provide evidence against a class (absence of negative evidence), Thus, our interest lies in the magnitude of *L*_SAM_,whose intensity should be increased to remain relevant to survival risk.

We conducted a comparison of SAM with various visualization methods from other published approaches.

Traditional attention-based heatmaps (TAH) illustrate the weights used for the average pooling of all WSI patches [12]. Co-attention heatmaps (CoAttn) display the co-attention weights assigned to each WSI patch and genomic signature [6]. TAH is applicable to models employing attention-weighted pooling of instances, while CoAttn suits multimodal models that fuse WSI and genomic features using co-attention. Consequently, we generated TAH results for ISM and MIL, and CoAttn results using GAM coattention weights for FSM. These were then quantitatively assessed against our SAMs.

## 3 Experiments

Using WSIs, bulk transcriptomics and survival data from three datasets (NLST, TCGA & CPTAC), we developed and validated an attention-based fusion framework by which to perform multimodal survival analysis. Our approach integrates bulk transcriptomics data with WSIs to predict patient survival, provides a means for topographic mapping of bulk gene expression on WSIs, generates WSI-level as well as an integrated multimodal spatial signature that points to tissue features associated with tumor survival on low- and high-risk cancer patients. We trained two models, one that used WSIs only (i.e., imaging survival model (ISM)), and the fusion survival model (FSM) that integrated WSIs and gene expression signatures. We used NLST for generating node-level features for graph construction [37], TCGA for training the models using 5-fold cross-validation, and CPTAC as an independent dataset for testing. We implemented the model using PyTorch (v1.12.1) and one NVIDIA 2080Ti graphics card with 11 GB memory on a GPU workstation. We set our model configurations as *L* = 3, *D* = 64 and *M* = 5, proportional to the number of gene signatures used in our paper. Considering that the imaging-graph has varying sizes, we used a batch size of 1. The training speed was about 5.1 iterations/s, and it took about 30 mins for each fold to reach convergence. The inference speed was 2.71 seconds per WSI with a batch size of 1.

### 3.1 Expert annotations

A subset of WSIs from the CPTAC cohort (10 cases) were uploaded to a secure, web-based software (PixelView; deepPath, Boston, MA). Using an Apple Pencil and an iPad, tumor regions were annotated by their histologic patterns (solid, micropapillary, cribiform, papillary, acinar, and lepidic). Tumor features including necrosis and vascular invasion were also annotated on the WSIs. Non-tumor regions were annotated as normal or premalignant airway epithelium, normal or inflammed lung, stroma, cartilage, and submucosal glands. We then evaluated the extent of overlap between the model-derived saliency maps and the expert-driven annotations.

### 3.2 Performance metrics

We reported cross-validated concordance index (c-index), which was averaged over the 5-folds. We also computed time-dependent area under the curve (tAUC) across 5-folds, which is a measure that evaluates how the model stratifies patient risk across various time points.

### 3.3 Comparison with prior work and ablation studies

#### 3.3.1 Model comparison with state-of-the-art approaches

We rigorously benchmarked our models, ISM and FSM, using the same 5-fold cross-validation framework as various state-of-the-art (SOTA) methods for survival prediction in computational pathology. Our evaluation is based on the most recent TCGA samples and excludes any samples without corresponding genomic data. For consistent comparison, we employed an identical SSL-based feature extraction process for WSIs and maintained uniform training hyperparameters and loss functions for all models. In cases where genomic data integration was required, we utilized the same genomic signatures identified in our models.

##### Unimodal baseline models versus ISM model

We adapted the survival neural network (SNN) architecture to utilize genomic features [15], specifically training it with our celltype gene signatures. To handle the diversity of gene signatures, we performed mean pooling across the groups before feeding them into a feed-forward network (FFN). The Attention MIL [12] method, a set-based neural network, employs global attention pooling to aggregate instance-level features, weighting each instance adaptively through a softmax function. DeepAttnMISL [35] starts by segmenting instance-level features into clusters via k-means, followed by a Siamese network processing for each cluster, and then aggregates the cluster features through global attention pooling. The Patch-GCN [4] approach treats WSIs as graphs, with patch features as nodes interconnected through k-NN, and derives the WSI representation by applying graph convolutional networks (GCNs) to this structure. TransMIL [21] and GTP [37] are transformer-based approaches. They aim to address the quadratic complexity issue, which arises due to the very large number of tokens (patches). TransMIL mitigates this complexity by replacing selfattention with the Nystrom method [32] in the transformer, while GTP [37] employs patch clustering using min-cut pooling [1] before applying the transformer. We standardized the node features for set-based methods like Attention MIL and DeepAttnMISL and used the same graph inputs for Patch-GCN as in our ISM model.

##### Multimodal baseline models versus FSM model

We compared the performance of various multimodal data fusion approaches with our FSM model. Specifically, we adapted PathomicFusion [5], which employs a region-of-interest (ROI) based approach, utilizing convolutional neural networks (CNNs) to extract features from H&E stained images, graph convolutional networks (GCNs) to analyze morphometric cell and graph features, and survival neural networks (SNNs) for genomic feature learning. These modalities are integrated for effective survival prediction. PORPOISE [3] utilizes Attention MIL for WSI feature interpretation and SSN for genomic insights, fusing these features for a comprehensive survival prognosis. MCAT [6] implements a co-attention mechanism to merge WSI and genomic features, which are then processed by a transformer for final survival outcome representation. MOTCAT [33] advances this integration by applying optimal transport to enhance global awareness and capture structural interactions within the tumor microenvironment for more accurate survival prediction. For each unimodal baseline, we enriched WSI features with genomic data; Attention MIL and DeepAttnMISL concatenate WSI features with genomic data for prediction, while Patch-GCN utilizes co-attention for feature fusion, aiming for improved predictive performance in survival analysis.

#### 3.3.2 Ablation studies

We conducted a series of ablation experiments to assess the impact of different feature extractors, node connectivity types, and the various components of the graph mixer layer (GML), specifically the node mixing layer (NML) and the channel mixing layer (CML). These ablation studies were carried out on both the ISM and FSM models to discern the contribution of each element. Furthermore, we evaluated the performance impact of the GML in the absence of the genomic module to understand its standalone efficacy.

### 3.4 Data and code availability

Data can be downloaded from the TCGA, CPTAC and NLST websites, respectively. The genomic data, python scripts and manuals are made available on GitHub (https://github.com/vkola-lab/tmi2024).

## 4 Results

Our graph attention-based framework demonstrated a robust ability to predict NSCLC survival. The ISM model’s performance exceeded that of other recent methodologies [4, 12, 21, 35, 37], as illustrated in Table 2. The ISM model demonstrated notable predictive performance on the TCGA dataset for both LUAD and LUSC cases. For LUAD cases, the ISM model achieved the highest c-index of 0.687 and for LUSC, it recorded 0.652. In terms of tAUC, the model showed its highest performance with values of 0.645 for LUAD and 0.647 for LUSC, highlighting its efficacy in predicting survival outcomes in these cancer types. For the CPTAC cohort, the ISM model’s performance was highest in LUSC with a c-index of 0.567 and a tAUC of 0.587 for LUAD. The TransMIL model slightly outperformed the ISM model in the CPTAC cohort for LUAD (c-index: 0.562 versus 0.540) and LUSC (tAUC: 0.661 versus 0.649). The ISM model’s performance highlights its efficacy in utilizing only WSIs, showcasing a competitive edge over other published methods that relied on genomic data or WSIs in isolation.

**Table 2:**
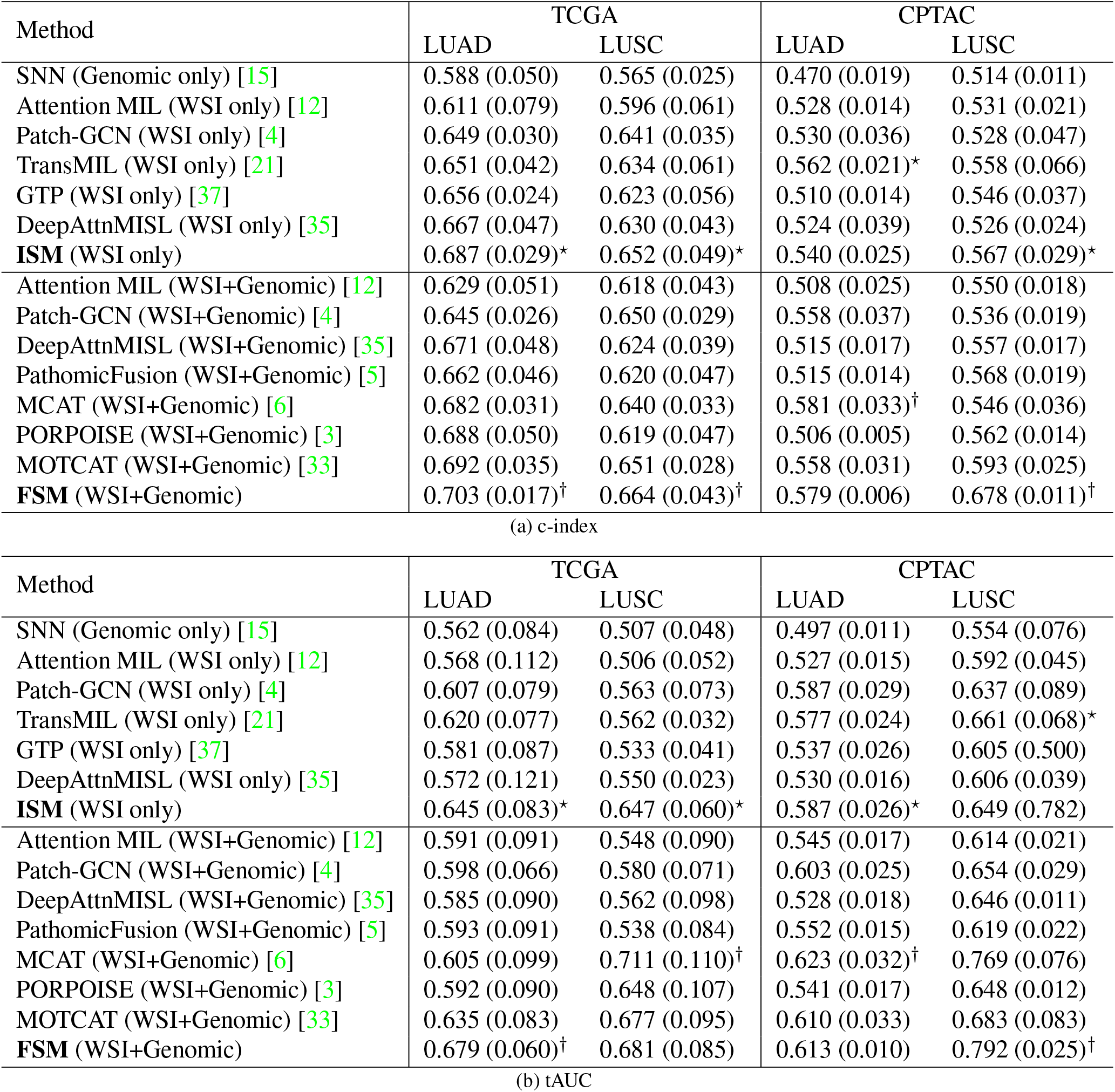
Model performance. Comparison of our models (ISM & FSM) with other published methods. The concordance index (c-index), and time-dependent area under the curve (tAUC) are shown. Five-fold cross validation was performed and mean as well as standard deviation (in parentheses) values are reported on the TCGA and CPTAC cohorts. The symbol *⋆* indicates the best c-index/tAUC values of the model that used WSIs, and the symbol † indicates the best c-index/tAUC values of the model that used WSI and genomic data. Text in bold indicates our models.

The FSM model that leveraged WSI and genomic data shows a marked improvement in performance across both metrics (c-index & tAUC) and datasets (TCGA & CPTAC), outperforming other methods such as Attention MIL [12], Patch-GCN [4], DeepAttnMISL [35], PathomicFusion [5], PORPOISE [3] and MOTCAT [33]. In the comparison between the FSM and MCAT [6] models across the TCGA and CPTAC cohorts, the FSM model consistently demonstrated superior performance in predicting survival outcomes for both LUAD and LUSC cases. Specifically, the FSM model achieved higher c-index values for LUAD and LUSC in the TCGA cohort (0.703 versus 0.682 and 0.664 versus 0.640, respectively), indicating a more accurate concordance in survival prediction. Although the FSM model’s c-index for LUAD was slightly lower in the CPTAC cohort (0.579 versus 0.581), it significantly outperformed the MCAT model for LUSC (0.678 versus 0.546), marking a notable advantage in predictive capability. Additionally, the tAUC values reinforced FSM’s robustness, particularly for LUAD cases on the TCGA cohort and LUSC cases on the CPTAC cohort, where FSM’s performance was distinctly better (0.679 versus 0.605 and 0.792 versus 0.769, respectively). These results highlight the FSM model’s potential to perform prognostic assessments in non-small cell lung cancer.

We observed a drop in the c-index values on the CPTAC cohort but the tAUC values were relatively similar to the TCGA cohort (Table 2). The reason for this disparity could be due to different sensitivities to time: tAUC is explicitly time-dependent and evaluates the model performance at various time points, whereas the c-index provides a general measure of concordance. If the model is highly sensitive to certain time intervals (performing well in those intervals and poorly elsewhere), this discrepancy could occur. The model was trained on TCGA whose range of survival time is [4, 7143] in days for LUAD, [0, 4765] in days for LUSC. The range of survival time in CPTAC is [0, 1836] days for LUAD, and [0, 1785] days for LUSC. So almost all CPTAC samples are high-risk cases with TCGA as the reference. The low c-index indicates that the ranking of all the high-risk cases is not favorable compared to TCGA. However, tAUC indicated that the model assesses the true positive and false positive rates well over various thresholds and time points.

From the ablation studies (Table 3), we observed: 1) The GML resulted in the best performance when NML and CML were included. NML is responsible for mixing information between different patches to learn local spatial relationships and fine-grained patterns within the WSI. CML is responsible for mixing information between channels to capture high-level interactions between features. Experiments shed light on the relative importance of NML versus CML, but both layers are essential to our approach. 2) Features based on semi-supervised learning (SSL) enhance model performance compared with ImageNet pretrained features. Experiments show that our model outperforms SOTA methods using ImageNet pretrained features to construct graphs. ResNet50 does not achieve better performance than ResNet18 because of the limited dataset (NLST) used for SSL. We observed that ResNet18 is sufficient and more efficient to achieve good performance. 3) NML using GAT performs better than NML using GCN. Experiments show that our model outperforms other SOTA methods using GCN as NML, highlighting the robustness of our approach. 4) We noticed that an 8-neighbor connectivity performs better than the 4-neighbor connectivity. An 8-neighbor connectivity considers diagonal as well as horizontal and vertical spatial connectivity between nodes. Important relationships between patches in the diagonal may be ignored when 4-neighbor connectivity is used. In the presence of noise or imperfections in an image, an 8-neighbor connectivity can provide better robustness as it incorporates more information from its neighbors.

**Table 3:** Ablation studies on model structures, graph featurization and graph construction. On our proposed ISM and FSM model architectures (rows 1 and 2, respectively), we performed ablation studies by removing or replacing several components of the architecture with other modules. We then compared the performance between these and the original ISM and FSM models. Conn: 4-node or 8-node connectivity in graph. Here GML: graph mixer layer, GCN: graph convolutional network, GAM: genomic attention module, CL: fine-tuning Resnet using contrastive learning on NLST, ImageNet: Resnet pretrained on ImageNet, c-index: concordance index, tAUC: time-dependent area under the curve, *⋆*: best performance on ISM model in each column, and †: best performance on FSM model in each column. Five-fold cross validation was performed on the TCGA cohort; mean and standard deviation values are reported. Of note, the FSM model refers to the cases with GAM and the ISM model refers to the cases without GAM.

| Graph Construction |  | Model |  | C-index |  | tAUC |  |
| --- | --- | --- | --- | --- | --- | --- | --- |
| Featurization | Conn | GML | GAM | LUAD | LUSC | LUAD | LUSC |
| CL Resnet18 | 8-node | NML+CML | x | 0.687 (0.029)* | 0.652 (0.049)* | 0.645 (0.083) | 0.647 (0.060)* |
| CL Resnet18 | 8-node | NML+CML | ✓ | 0.703 (0.017)† | 0.664 (0.043)† | 0.679 (0.060)† | 0.681 (0.085)† |
| ImageNet Resnet18 | 8-node | NML+CML | ✓ | 0.661 (0.055) | 0.638 (0.022) | 0.581 (0.089) | 0.619 (0.033) |
| CL Resnet18 | 8-node | x | x | 0.589 (0.022) | 0.518 (0.014) | 0.552 (0.072) | 0.554 (0.043) |
| CL Resnet18 | 8-node | NML | x | 0.654 (0.021) | 0.607 (0.030) | 0.604 (0.072) | 0.646 (0.071) |
| CL Resnet18 | 8-node | CML | x | 0.589 (0.022) | 0.532 (0.025) | 0.567 (0.083) | 0.565 (0.033) |
| CL Resnet18 | 8-node | x | ✓ | 0.588 (0.042) | 0.540 (0.025) | 0.602 (0.044) | 0.555 (0.036) |
| CL Resnet18 | 8-node | NML | ✓ | 0.667 (0.028) | 0.622 (0.016) | 0.628 (0.050) | 0.658 (0.034) |
| CL Resnet18 | 8-node | CML | ✓ | 0.593 (0.032) | 0.541 (0.024) | 0.623 (0.069) | 0.642 (0.029) |
| ImageNet Resnet18 | 8-node | x | x | 0.549 (0.048) | 0.529 (0.018) | 0.601 (0.050) | 0.561 (0.027) |
| ImageNet Resnet18 | 8-node | NML+CML | x | 0.658 (0.043) | 0.619 (0.033) | 0.624 (0.061) | 0.632 (0.021) |
| CL Resnet50 | 8-node | NML+CML | x | 0.679 (0.040) | 0.642 (0.027) | 0.675 (0.053)* | 0.644 (0.041) |
| CL Resnet18 | 8-node | GCN+CML | x | 0.677 (0.044) | 0.598 (0.040) | 0.630 (0.064) | 0.623 (0.043) |
| CL Resnet18 | 8-node | GCN+CML | ✓ | 0.685 (0.043) | 0.647 (0.030) | 0.663 (0.057) | 0.667 (0.033) |
| Imagenet Resnet18 | 8-node | GCN+CML | x | 0.622 (0.012) | 0.602 (0.037) | 0.605 (0.037) | 0.634 (0.039) |
| Imagenet Resnet18 | 8-node | GCN+CML | ✓ | 0.637 (0.023) | 0.623 (0.045) | 0.644 (0.029) | 0.667 (0.025) |
| CL Resnet18 | 4-node | NML+CML | x | 0.667 (0.023) | 0.638 (0.038) | 0.644 (0.066) | 0.622 (0.061) |
| CL Resnet18 | 4-node | NML+CML | ✓ | 0.691 (0.037) | 0.648 (0.034) | 0.649 (0.065) | 0.670 (0.032) |

Our framework is also capable of generating interpretable maps which compare favorably with expert-driven annotations. The generated SAMs pointed to WSI regions that were associated with prognostic histologic features and patterns (Fig. 4). Qualitatively, we observed a high degree of overlap between the pathologist tumor region annotations with the salient tissue regions identified by the SAMs. For example, the LUAD high- and low-risk cases and the LUSC high-risk case all have non-tumor tissue that is not highlighted by the models. The FSM model, compared to ISM, more strongly highlighted tumor histologic patterns and features that have known associations with prognosis. For example, in the LUAD low-risk case, the FSM model more strongly highlighted the lepidic tumor histologic pattern compared to the more aggressive solid pattern [11, 14, 18]. In the LUAD high-risk case, the FSM model highlighted the aggressive solid tumor histologic pattern and the focus of vascular invasion [24, 34] associated with an unfavorable prognosis. In LUSC, there are a few known histologic patterns or features associated with prognosis, however, we observe that in the low-risk LUSC case, the model highlighted tumor regions with high immune infiltrate that may be important to the survival prediction. Interestingly, the SAMs for both the ISM and FSM models localized similar neighborhoods as highly associated with patient survival, with FSM model often highlighting additional tumor-specific regions.

**Figure 4.**
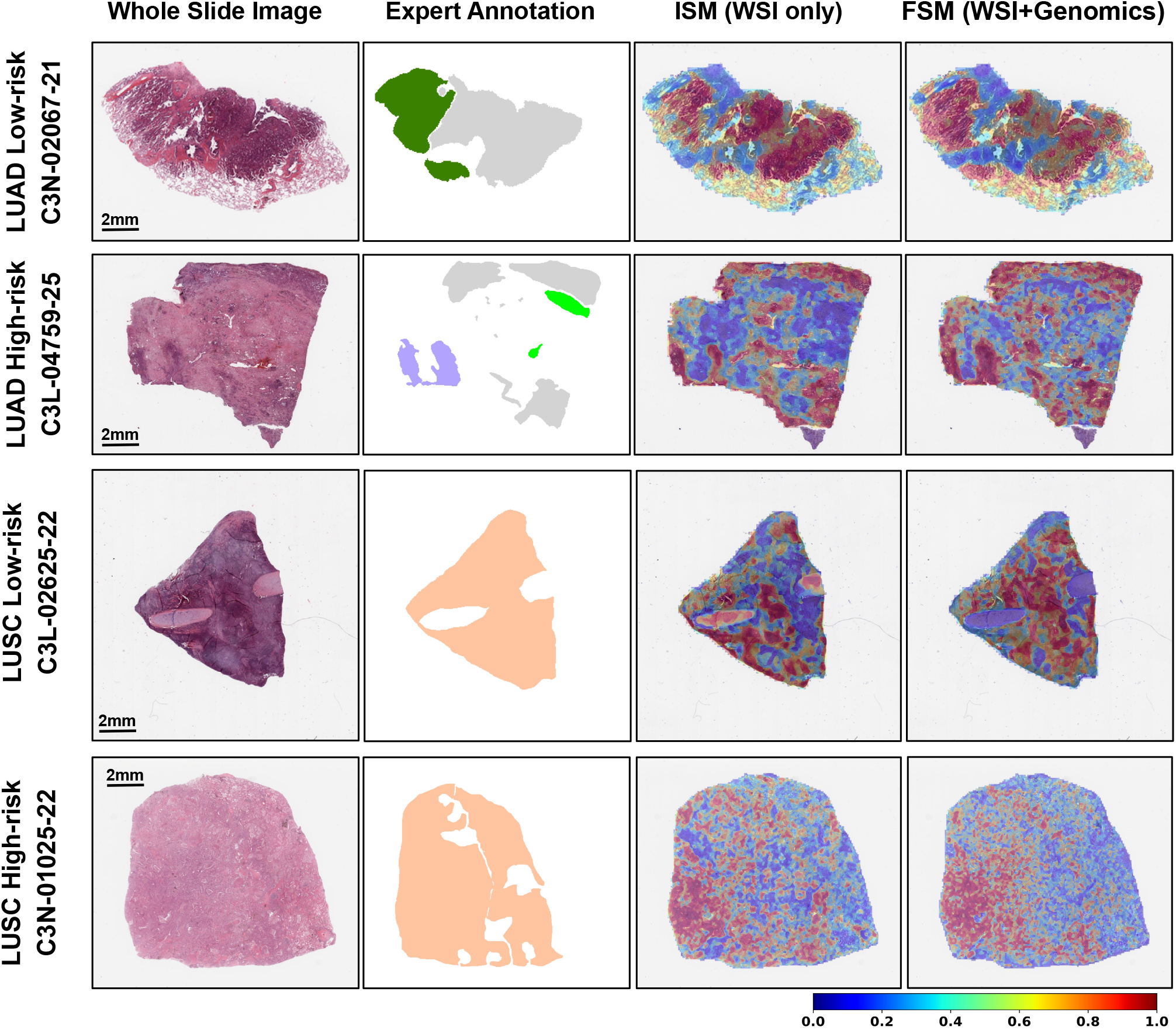
Survival activation map (SAM) on human NSCLC samples. The first column shows the H&E WSIs, the second column shows the pathologist annotations of the tissue, the third, and fourth columns indicate the SAMs based on the ISM and the FSM models, respectively. Top row, low-risk LUAD case where annotations are: low-risk related lepidic (dark green) and high-risk related tumor (light gray) histologic patterns. Second row, high-risk LUAD case where annotations are: high-risk related solid histologic pattern (lavender), high-risk related vascular invasion (light green), and low-risk related tumor (light gray) histologic patterns. Third and fourth rows, low-risk and high-risk LUSC cases, respectively where annotations are: tumor tissue (peach). The colorbar is relevant to the heatmaps shown in the last two columns.

We introduced additional visualizations to benchmark our SAM framework against traditional attention-based heatmap (TAH) applied to the ISM model and on Attention-MIL [12], which is a multiple instance learning framework (MIL) (Fig. 5). The TAH visualizes the weights assigned to nodes, and they are considered as the ‘importance’ of nodes after MIL is trained. The softmax function in TAH is sensitive to large values in the node’s weights. Large values can lead to extremely small attention for the other nodes, which may not reflect the actual uncertainty or variability in the data. In comparison, our approach uses both the gradients and activations within the graph network, and this provides a balance between node details (from activations) and semantic information (from gradients). As we used co-attention to integrate WSI patches and gene signatures, we also generated visualizations based on the attention scores assigned to each patch for each gene signature (denoted as CoAttn). We then computed Dice coefficients [23] to quantitatively assess the similarity between pathologist annotations and the salient tissue regions identified by SAM, TAH, and CoAttn for different models (Fig. 6). The FSM model’s SAM, which integrates cell-type signatures, aligns closely with pathologist annotations, indicating its clinical utility. Co-Attention (CoAttn) visualizations demonstrate the interaction of WSI patches with genomic signatures, but SAM directly associates these regions with survival outcomes. Higher Dice coefficients across various thresholds confirm SAM’s enhanced performance, as it captures all gene expression signatures and identifies prognostic areas within the WSI. CoAttn visualizations, while occasionally coinciding with pathologic annotations, do not consistently match the SAM’s comprehensive coverage, which includes all gene expression signatures for identifying survival-associated regions.

**Figure 5.**
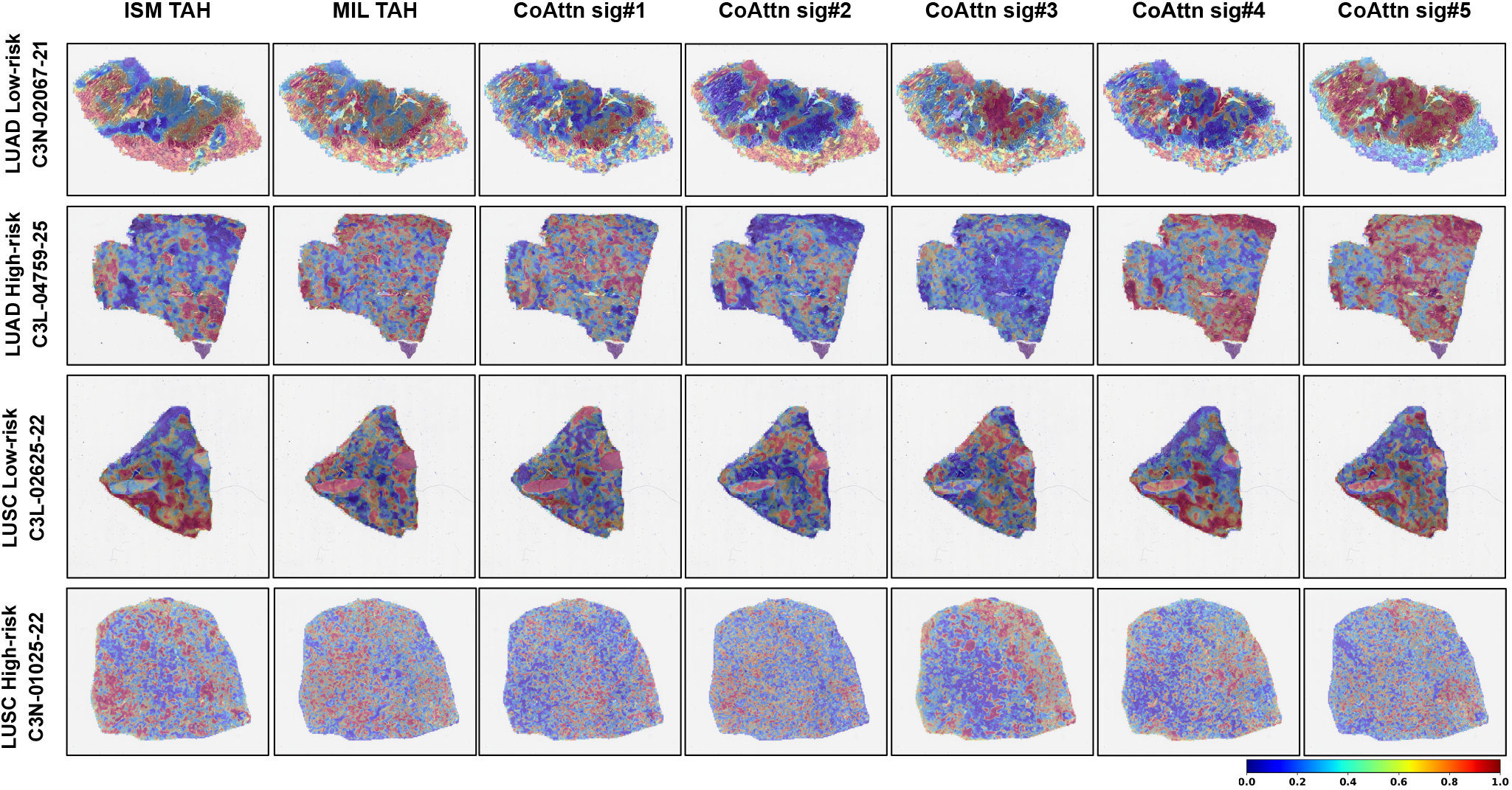
Visualization of salient regions using various interpretable methods. WSI-level heatmaps highlighting WSI regions associated with survival on four (low-and high-risk LUAD as well as low-and high-risk LUSC) cases are shown (see Fig. 4 for more info). The first column shows the traditional attention-based heatmaps (TAH) generated on the ISM model and the second column shows the ones generated on a multiple instance learning model (Attention MIL [12]). The remaining columns show the co-attention (CoAttn) heatmaps generated on the FSM model for the gene signatures, sig#1-sig#5 (see Section 2.1), respectively.

**Figure 6.**
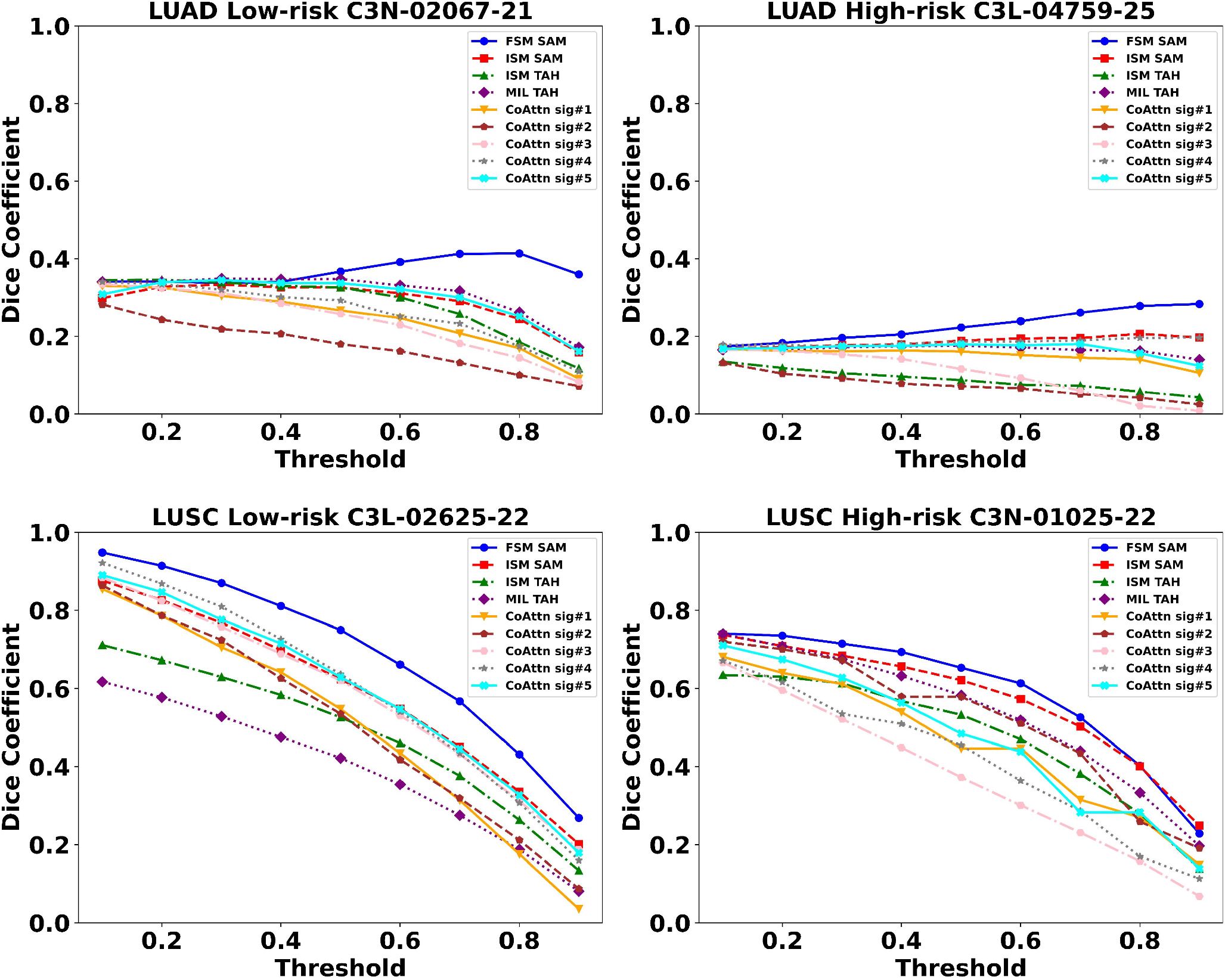
Quantitative comparison of different model interpretability methods with expert annotations. The plots display the performance of the FSM SAM (blue), ISM SAM (red), ISM TAH (green), MIL TAH (purple), CoAttn sig#1 (orange), CoAttn sig#2 (brown), CoAttn sig#3 (pink), CoAttn sig#4 (gray), and CoAttn sig#5 (cyan) interpretability methods, in terms of the Dice coefficient across different threshold levels. For each case, the Dice coefficient was computed by generating binarized heatmaps at different thresholds, and comparing them with the pathologist annotations. ISM TAH, MIL TAH, and CoAttn heatmaps are shown in Fig. 5. The four cases correspond to the low- and high-risk LUAD as well as the low- and high-risk LUSC cases presented in Fig. 4, which also include the pathologist annotations and the SAMs. In the LUAD cases, the light gray annotations indicating tumor regions were excluded from the Dice coefficient calculation. This exclusion is because we focused solely on pathologic tumor features or patterns known to be associated with either favorable prognosis in low-risk LUAD or unfavorable prognosis in high-risk LUAD.

## 5 Discussion

Our interpretable deep learning approach can perform attention-based fusion of WSIs and bulk transcriptomics data to predict NSCLC survival. By the standards of various metrics, our approach displayed superior performance compared with the SOTA approaches, yielding consistent predictions on two different sample sets ± TCGA and CPTAC. Beyond model performance, we can generate attention-based SAMs that highlight regions on the WSIs that correspond to prognostic tumor histologic patterns and features identified via expert annotations on low- and high-risk NSCLC cases. Additionally, the SAMs identified WSI regions that extended beyond the tumor regions to reveal image-genomic relationships that could be implicated in patient survival.

The attention mechanism serves to enhance model performance by focusing on the most relevant aspects of each data modality in a context-aware manner. Additionally, our framework aids in capturing complex interdependencies between images and gene expression at varying levels of granularity. Another significant technical advantage is the interpretability of the model’s decision-making process - the attention-based mechanism can highlight the important features in each modality, providing valuable insights into the model’s rationale. The graph attention layer enabled every node in the imaging graph attend to its neighbors given its own representation as the query so that the local relationships are better learned than the previously published methods. Furthermore, by zeroing in on the salient data sections in each modality, our framework boosts computational efficiency, reducing the processing load without compromising on the model performance.

In our study, we compared model-based saliency maps with expert-driven annotations on a small set of cases and thus our conclusions are limited. The small set of cases was selected because manual annotation is a tedious task, and the pathologist’s availability was limited. Moreover, the pathologist annotated histologic patterns and features, some of which are associated with survival, but a larger study that includes pathologic annotation of tumor tissues and spatial omics is needed to evaluate the regions highlighted by both the ISM and FSM models. In addition, the negative log-likelihood (NLL) loss function used in our model has some limitations that include the assumption that the proportional hazards is integral to the likelihood formulation. If this assumption is violated (i.e., the hazard ratios are not constant over time), the NLL optimization may produce biased estimates. Censored observations can also complicate the estimation process as they provide partial information about the survival time. Too much censoring can lead to imprecise estimates, affecting the robustness of the optimization. The censoring bias in survival prediction presents a significant challenge to model training, especially as new datasets may exhibit widely varying censoring rates. To handle the censoring effect in survival analysis, future work could use the inverse probability of censoring weighting to create a pseudo-population that is representative of the population without censoring. By re-weighting individuals based on their probability of being uncensored, one can potentially reduce bias due to censoring.

In conclusion, our graph attention-based approach can efficiently process WSI and bulk genomic data and estimate NSCLC survival. Future work will include testing our model using various cell type and prognostic gene expression signatures that are implicated in survival of various cancers. Additional studies to generate data on NSCLC specimens using modern spatial technologies will help validate the biological insights obtained via SAMs. Extension of this framework to other cancers and various types of omic data is needed to fully appreciate its broad potential in performing multimodal survival analysis.

